# Estimating colocalization probability from limited summary statistics

**DOI:** 10.1101/2020.05.19.104927

**Authors:** Emily A. King, Fengjiao Dunbar, Justin Wade Davis, Jacob F. Degner

## Abstract

1

**Motivation:** A common approach to understanding the mechanisms of noncoding GWAS associations is to test the GWAS variant for association with lower level cellular phenotypes such as gene expression. However, significant association to gene expression will often arise from linkage disequilibrium to a separate causal variant and be unrelated to the mechanism underlying the GWAS association. Colocalization is a statistical genetic method used to determine whether the same variant is causal for multiple phenotypes and is stronger evidence for understanding mechanism than shared significance. Current colocalization methods require full summary statistics for both traits, limiting their use with the majority of reported GWAS associations (e.g. GWAS Catalog). We propose a new approximation to the popular coloc method [1] that can be applied when limited summary statistics are available, as in the common scenario where a GWAS catalog hit would be tested for colocalization with a GTEx eQTL. Our method (POint EstiMation of Colocalization - POEMColoc) imputes missing summary statistics using LD structure in a reference panel, and performs colocalization between the imputed statistics and full summary statistics for a second trait.

**Results:** As a test of whether we are able to approximate the posterior probability of colocalization, we apply our method to colocalization of UK Biobank phenotypes and GTEx eQTL. We show good correlation between posterior probabilities of colocalization computed from imputed and observed UK Biobank summary statistics. We perform simulations and show that the POEMColoc method can identify shared causality with similar accuracy to the coloc method. We evaluate scenarios that might reduce POEMColoc performance and show that multiple independent causal variants in a region and imputation from a limited subset of typed variants have a larger effect while mismatched ancestry in the reference panel has a modest effect.

We apply POEMColoc to estimate colocalization of GWAS Catalog entries and GTEx eQTL. We find evidence for colocalization of ~ 150,000 trait-gene-tissue triplets. We find that colocalized trait-gene pairs are enriched in tissues relevant to the etiology of the disease (e.g., thyroid eQTLs are enriched in colocalized hypothyroidism GWAS signals). Further, we find that colocalized trait-gene pairs are enriched in approved drug target - indication pairs.

**Availability:** POEMColoc is freely available as an R package at https://github.com/AbbVie-ComputationalGenomics/POEMColoc

## 2 Introduction

Genome-wide association studies have identified thousands of disease and trait associated loci but as most GWAS associated loci are non-coding, their functional interpretation remains difficult. eQTL studies are a popular way to follow up on GWAS studies. By providing a link between an associated variant and a gene, they facilitate functional interpretation of GWAS results, especially when associated variants are found in noncoding regions of the genome. A simple approach is to query eQTL datasets for GWAS lead variants to determine if they are significant eQTL for any gene. However, this approach can lead to false positive GWAS-eQTL links as linkage disequilibrium can lead to shared significance at a SNP without the two traits sharing a causal variant. Colocalization analysis reduces false positive results by directly testing competing hypotheses of causal sharing.

Regulatory Trait Concordance (RTC) [2] is early method for estimating causal sharing that may be applied to situations in which only a GWAS top SNP is known. However, it requires individual-level data for the second dataset (*e.g*. eQTL) and does not actually provide a probability of causal sharing, merely a score that can be used to prioritize the most likely causal links. Coloc [1] and its multi-trait extension moloc [3] are popular colocalization methods with an efficient and easy to use R implementation. They require full summary statistics for both traits and compute the probability of each causal hypothesis using approximate Bayes factors. eCAVIAR [4] is another colocalization method that can also be applied to summary statistics (supplemented with LD information) that has the additional advantage of being able account for more than one causal variant in a region. Enloc [5] is a third approach to colocalization using a Bayesian hierarchical model to compute a regional colocalization probability within an LD block containing a GWAS signal.

One limitation for using most of these colocalization methods is that full summary statistics for GWAS studies are frequently not available. For example, the largest repository of human trait associated variants, the GWAS catalog, only reports statistics for the top-associated variants in a given study. For this reason, until recently, only simple approaches such as checking the eQTL significance of the reported GWAS variant were possible. Like the POEMColoc method, PICCOLO [6] was recently developed to compute colocalization of signals when only the top SNP is available. However, POEMColoc takes a different approach that does not discard information when full summary statistics are available for one but not both of the traits and does not assume that both traits have a causal variant so that probabilities of all coloc hypotheses are computed.

## 3 Methods

### 3.1 Method Overview

Here, we propose a method POEMColoc (POint EstiMation of Colocalization) where the explicit goal is to approximate the coloc method [1] under the commonly encountered constraint wherein full summary statistics *Z*_1,1_,…,*Z*_1,*p*_ are available for one trait at *p* SNPs (*e.g*, Heart Atrial Appendage expression of SCN5A) while summary statistics for only the top SNP *Z*_2,*obs*_ in a region is reported for the second trait (*e.g*. a SNP associated with QT interval from the GWAS catalog). Like the original coloc method, we compute posterior probabilities of five mutually exclusive and exhaustive hypotheses:

- *H*_0_: Neither trait has a causal SNP in the region
- *H*_1_: Only trait 1 has a causal SNP in the region
- *H*_2_: Only trait 2 has a causal SNP in the region
- *H*_3_: Both traits have a causal SNP in the region, but the two causal SNPs are different
- *H*_4_: Both traits have a causal SNP in the region, and the two causal SNPs are the same of which, we are primarily interested in the posterior probability of *H*_4_.

Our method first imputes missing summary statistics using both the summary statistics for the reported variant and the LD structure of the region in a reference population (Figure 1). For a single causal SNP indexed by *c*, Z scores at the full set of SNPs can be approximated by a multivariate normal distribution with parameters

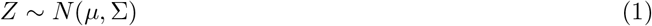

where

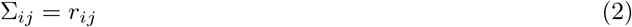

is the Pearson correlation between SNPs *i* and *j* and

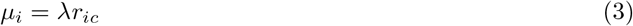

where *r_ic_* is the Pearson correlation between SNP *i* and the causal SNP *c* [7]. A reasonable point estimate of the unknown parameters is λ = *Z_obs_* and *c* is the index of the causal SNP (in fact, this estimate is a maximizer of the full data likelihood under the approximate normal distribution, Supplement Text S1). Using this point estimate of λ and *c*, we can compute the expected value of the missing summary statistics conditional on the observed summary statistic at the top SNP. This conditional expectation formula has been used in previous work on the imputation of summary statistics at untyped SNPs [8][9].

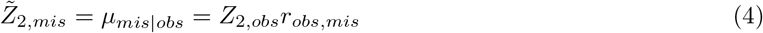

**Figure 1:**
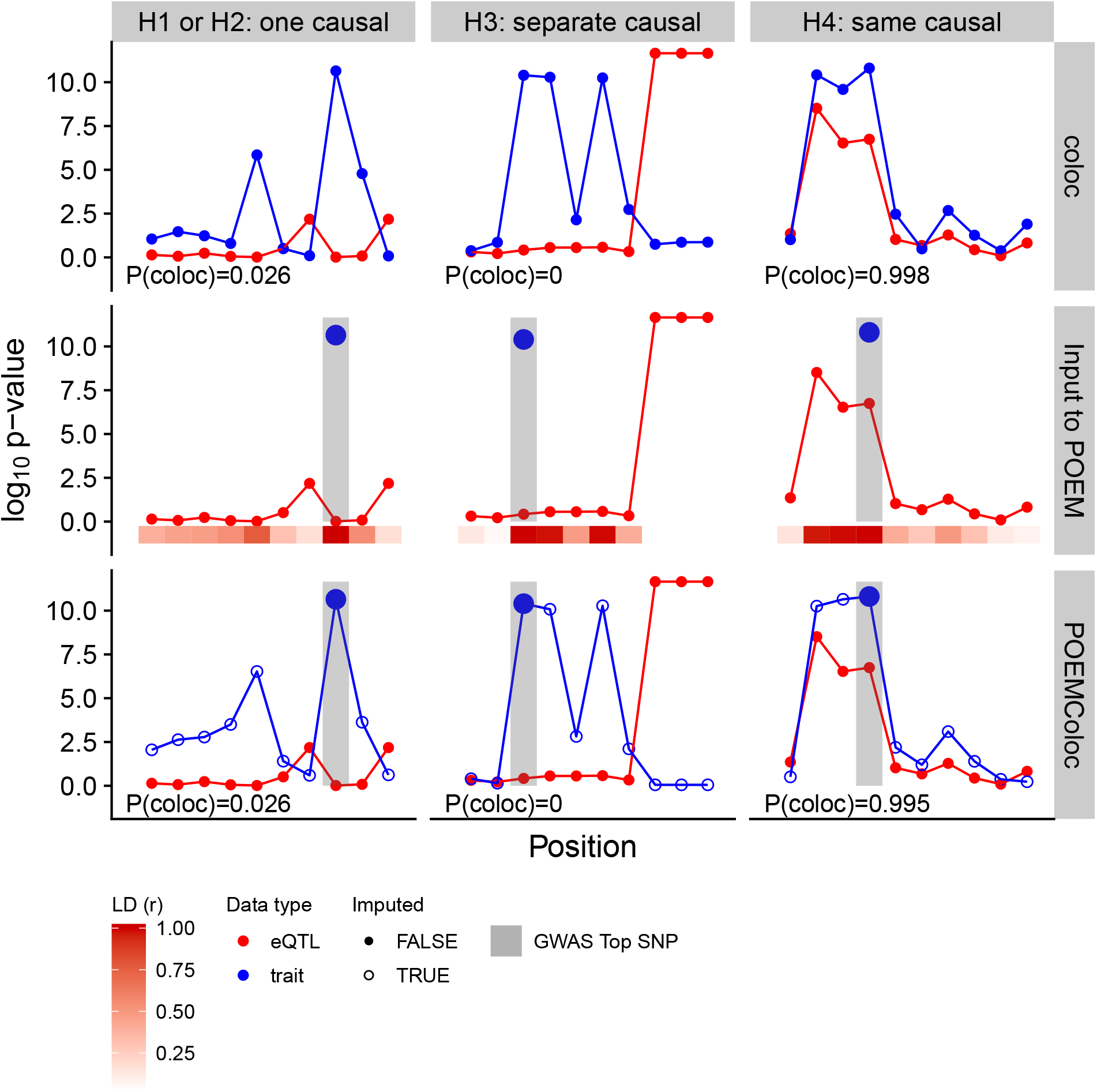
Illustration of the POEMColoc method compared to coloc. Under *H*_1_ or *H*_2_, only one of the two traits has any causal SNP in the region. Under *H*_3_, the two traits have two different causal SNPs. Under *H*_4_ (colocalization) the two traits share a causal SNP. The first row of the plot illustrates the coloc method using *p*-values. Input data is *p*-value at each SNP in the region for each trait, and the output is a posterior probability of each hypothesis. The posterior probability of colocalization (*H*_4_) is shown on the plot. The second row illustrates input to the POEMColoc method. Full summary statistics are available for one trait only, and for the second trait only the position and *p*-value of the top SNP is known. We also require LD from the top SNP to at least some of the trait 1 SNPs to be known from a reference panel (shown below in red). We use the LD to impute missing *p*-values for input to coloc. Imputed *p*-values are show in the bottom row. POEMColoc consists of applying coloc to the imputed datasets, outputting posterior probabilities of colocalization show in the bottom row.

Similar to coloc, *Z* scores used in POEMColoc may either be computed from *p*-values or from coefficient estimates *β* and var*β*. In practice, we expect *p*-values to be the most common form of input to POEMColoc. We transform two-sided *p*-values to Z-scores using inverse cdf function *F Z* = *F*^−1^(*p*/2) We then apply *F* to the imputed values 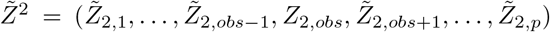 to obtain two-sided *p*-values *F*(-|*Z*|)/2 and use them as input to `coloc.abf` from the `coloc R` package, along with *Z*_1_. In many cases, such as the UK Biobank association statistics we analyze in this paper, *p*-values are known to have been generated from a *t*-distribution with approximately *N* degrees of freedom, but in the absence of other information we may also take *F* to be the standard normal cdf. When neither of the datasets have full summary statistics, the imputation is performed for both datasets on all SNPs in a reference panel within a user-specified window of either top SNP, optionally subject to a minor allele frequency cutoff. When necessary, we will distinguish these two versions of coloc using POEMColoc-1 (POEMColoc with one imputed dataset) and POEMColoc-2 (POEMColoc with two imputed datasets). Like the *p*-value implementation of coloc, POEMColoc additionally requires sample size and, for case-control studies, case fraction. Minor allele frequency and *r* are either user-supplied or computed from a reference panel using SeqArray [10].

### 3.2 Evaluating POEMColoc on UKBB GWAS hits and GTEx eQTL

Full summary statistics for GWAS conducted on UKBB phenotypes were downloaded on 4 June 2019 from http://www.nealelab.is/uk-biobank/. We removed duplicated phenotypes by excluding sex-specific GWAS runs and excluding raw quantitative variables in favor of the inverse rank transformed alternative. For each GWAS, we selected non-overlapping windows around genome-wide significant (*P* < 5 × 10^-8^) lead SNPs as candidates for colocalization with eQTL. Variants marked as low confidence were excluded.

As a test case to evaluate the performance of the POEMColoc method, for each UKBB window described above, we selected eQTL summary statistics from GTEx release 7 whole blood eQTL of the entire cis-candidate window of any gene containing associated UKBB variant within its cis-candidate window [11]. From the 2197 UKBB phenotypes with at least one genome-wide significant hit, there were 1553 phenotypes where at least one of the genome-wide significant hits overlapped a tested eQTL in GTEx whole blood. We merged summary statistics for all variants in common between the GTEx eQTL tests and the UKBB phenotype tests and used each gene, UKBB phenotype, and UKBB selected region as a way to evaluate POEMColoc in comparison to coloc.

For each combination of UKBB and GTEx summary statistics, we applied coloc via the coloc.abf function using default priors. On the same combinations of UKBB and GTEx summary statistics, we ran POEMColoc using the full list of *p*-values from GTEx eQTL but using only the top SNP from the UKBB region as if it was reported in the GWAS catalog. We used LD information from unrelated European individuals in the 1000 Genomes phase3 reference panel as processed/distributed for use with the BEAGLE imputation software package (http://bochet.gcc.biostat.washington.edu/beagle/1000_Genomes_phase3_v5a/). We evaluated performance of POEMColoc under the scenario where the reported GWAS SNP came from a limited SNP subset by choosing the UKBB top SNP from only those SNPs contained on each of three Illumina genotyping chips that vary in total SNP content from 1 million to 5 million SNPs (1M - OmniExpressExom8; 2.5M - HumanOmni2.5Exome; 5M - HumanOmni5Exome).

### 3.3 Evaluating POEMColoc performance with simulations

Simulation studies were performed using genotype data from UK Biobank British ancestry individuals and simulated phenotypes. The UK Biobank was approved by the North West Multi-centre Research Ethics Committee. This research was conducted under UK Biobank resource application 26041. For each simulated dataset, a causal eQTL SNP was randomly selected among variants with minor allele frequency at least 0.01. Under H3, a second, distinct causal SNP was randomly selected from variants within a 1 kb window from the first causal GWAS SNP. Close proximity of the two causal SNPs ensures that many datasets will present a challenging colocalization problem, in which the causal GWAS SNP may have a statistically significant association without shared causality due to linkage disequilibrium. We randomly sampled *N*_1_ = 500 individuals to simulate eQTL phenotypes, and *N*_2_ = 10000 new individuals to simulate GWAS phenotypes. The phenotype for *i*th individual in group *k*(*Y_ki_*) is simulated based on the genotype from UKBB data, 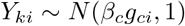 where 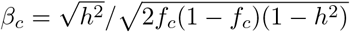 is computed from the minor allele frequency *f_c_* at site *c* and single SNP heritability *h*^2^. *h*^2^ values were sampled from beta distributions with means 0.0099 for GWAS and 0.1 for eQTL (Beta(4, 400) and Beta(2, 18) respectively). We compared POEMColoc with coloc, PICCOLO and with the simple approach using top SNP eQTL *p*-values. As in other analyses, coloc and POEMColoc were run using the default coloc priors on causal configurations.

### 3.4 Comparison to PICCOLO

POEMColoc was compared to PICCOLO using its default settings and PICS causal probabilities obtained from pics.download using the EUR ancestry option. We attempted to run PICCOLO on all simulated datasets and a random sample of 100 UK Biobank phenotypes with at least one genome-wide significant association. Due to its use of PICS tool for estimating causal SNP probabilities, PICCOLO is not able to run on a large fraction of analyzed datasets (Table S1). All comparisons presented between PICCOLO and POEMColoc are restricted to the subset of datasets on which both could be run.

### 3.5 Running POEMColoc on GWAS catalog

GWAS Catalog data were downloaded on Nov 26, 2018. For each GWAS catalog entry, we attempted to obtain the SNP *p*-value, whether the study is case-control or quantitative trait, the sample size *N*, and, for case-control studies, the fraction of observations that are cases. We also attempted to assign each entry to a broad ancestry group in order to choose an appropriate reference panel for calculation. We excluded associations for which case-control or quantitative trait status and sample size could not be ascertained using our automated approach, and those not meeting the *p*-value threshold 5 × 10^-8^. For each GWAS Catalog SNP rsid, we extracted GTEx summary statistics for all genes and tissues with available summary statistics overlapping the GWAS SNP. When a matched ancestry panel was available, POEMColoc was implemented using both the matched ancestry panel and the full 1000 Genomes from all unrelated individuals; otherwise the full 1000 Genomes was used. Using those associations for which a matched reference panel was available, we show that more colocalizations were detected using a matched reference (Figure S4). For subsequent analyses, we used colocalization results from the matched ancestry panel where available and otherwise used the full 1000 Genomes. We evaluated the biological relevance of GWAS Catalog colocalizations using two different metrics, tissue enrichments of eQTL signals, and enrichments for approved drug mechanisms.

#### 3.5.1 Estimating tissue enrichment of eQTL signal

We collapsed replicate associations to obtain one colocalization posterior probability per gene-trait-tissue trio as the maximum across associations. Using a colocalization cutoff of 0.9, we determined an enrichment score for each trait-tissue combination as the −log_10_ *p*-value from Fisher’s exact test for enrichment of colocalizations in the tissue-trait pair (Supplement Text S2).

#### 3.5.2 Association with approved drug targets

We used target-indication pair approval status from the supplementary materials of [12] to assess whether colocalization was associated with approval. xMHC targets (Chromosome 6 25.7 Mb-33.4 Mb) are unavailable because they were excluded from this analysis. Colocalizations were collapsed at the level of gene-trait pair by taking the maximum probability across tissues and genes. For each gene-trait pair assessed for colocalization, we determined whether or not it matched an approved drug mechanism. eQTL evidence classes were determined strong evidence for colocalization (*P*(*H*_4_) ≥ 0.9) evidence against colocalization (*P*(*H*_4_) ≤ 0.5) and significant eQTL for the GWAS top SNP (*p* < 10^-6^), and combinations thereof. We computed the odds ratio of such a match for different positive classes of eQTL evidence relative to candidate pairs with no evidence of an eQTL via colocalization or eQTL *p*-value. It was determined that different evidence classes differed systematically in the proportion of coding genes and observed drug targets, so we conditioned this analysis on the candidate gene being an approved drug target. Significance was assessed via permutation of GWAS trait labels, which further helps separate ubiquitous target-level variation in colocalization and eQTL probability from the effect of the match between the target and the indication.

#### 3.5.3 PubMed Odds Ratio similarity

In order to identify drug mechanisms supported by colocalizations and to have a quantitative assessment of tissue-trait similarity, we used similarity in the MeSH vocabulary. Similarity is determined through odds ratio of cooccurance in article MeSH terms in the PubMed corpus (accessed October 10, 2018), an approach based on [13]. For identifying similar drug mechanisms, we used an odds ratio cutoff of 20 as this corresponded on average to the similarity cutoff used in the previous work. Drug indications are provided as MeSH terms in the supplementary materials of[12] and GWAS traits and tissues were mapped to the MeSH vocabulary using a similar procedure. Because of concerns about circularity from, for example, an approved drug leading to occurrence of its target and indication in publication, we exclude MeSH terms under the headings Amino Acids, Peptides, Proteins, Enzymes and Coenzymes, and Genetic Phenomena as well as all studies related to drug response or adverse reactions from the set of GWAS traits analyzed.

## 4 Results

### 4.1 Approximation accuracy using UK Biobank phenotypic associations and GTEx expression QTL

To assess the accuracy of our method at recovering the colocalization posterior probabilities using full summary statistics and the original coloc method, we used the combination of quantitative trait associations from the UK Biobank (UKBB) and whole blood gene expression associations from the GTEx project [11].

In total, we obtained 646884 combinations of UKBB phenotype associated window and GTEx eQTL summary statistics. We ran colocalization analysis on each of these using the original coloc method with full summary statistics. We find that approximately 0.4% of tested pairs are colocalized (*P*(*H*_4_) > 0.9).

Next, we imagined the scenario in which instead of having full summary statistics for the UKBB phenotypes, these phenotypes had been reported in biomedical literature and only the most significantly associated variant for each locus was available. We used our method as implemented in the associated R package POEMColoc. Posterior probabilities of each hypothesis were highly correlated between POEMColoc and coloc (Figure 2A, *R*^2^ = 0.81 for *P*(*H*_4_)). Use of *p*-values in place of regression coefficients as the input to coloc improves correlation for some hypotheses but has minimal effect on the posterior probability of colocalization *H*_4_ (Figure S1). Furthermore, treating colocalization as a binary classification problem where full coloc *P*(*H*_4_) > 0.9 was considered a true positive and all other outcomes were considered true negatives, we achieve a high degree of sensitivity and specificity using POEMColoc (AUCPR=0.91 for POEMColoc-1 and 0.84 for POEMColoc-2). We performed colocalization analysis for a subset of phenotypes using PICCOLO, and found POEMColoc’s performance to more closely match that of coloc, even when using only data from the top SNP from both traits (Figure 2B, Table S1, Figure S2).

**Figure 2:**
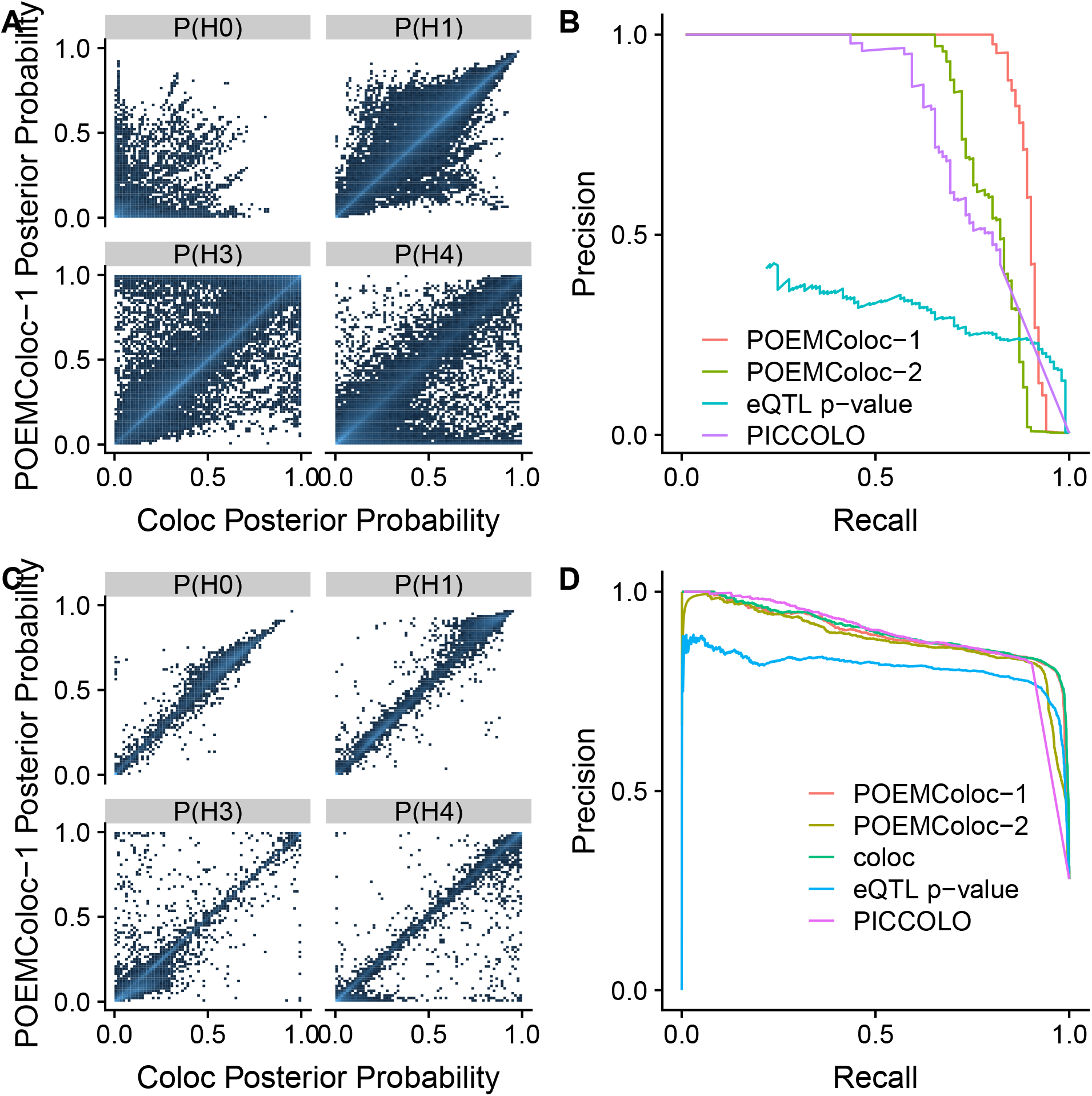
A: Comparison of hypothesis posterior probability using POEMColoc-1 method to coloc using GTEx eQTL regression coefficients and UK Biobank *p*-value summary statistics as input B: Precision-recall curves for predicting colocalization using coloc (*P*(*H*_4_) > 0.9) in the UK Biobank. C: Comparison of hypothesis posterior probability between POEMColoc-1 and coloc in simulated datasets. D: Precision-recall curves for predicting colocalization (*H*_4_) in simulated datasets, compared to coloc and a simple method using the eQTL *p*-value of the GTEx top SNP.

### 4.2 Methods comparison for simulated associations

While our stated goal was to approximate coloc using full summary statistics, performance assessments using coloc as the gold standard cannot compare performance between coloc and POEMColoc at correctly assigning associations to the true hypothesis. To address these limitations we compared the performance of POEMColoc and coloc on simulated data and compared both to PICCOLO and to what performance would be if we used the simple approach of quantifying causal evidence using the eQTL *p*-value of the top GWAS SNP. Hypothesis posterior probabilities calculated with POEMColoc are highly correlated with posterior probability calculated with coloc method (Figure 3C, *R*^2^ =0.98 for the posterior probability of colocalization). The precision recall plot in Figure 3D shows that the performance of POEMColoc is close to coloc method, better than the simple method (eQTL value) of using the eQTL *p*-value of the GWAS top SNP, and has higher AUCPR than PICCOLO, even when using only the top SNP from each dataset (Table S1). A further advantage of POEMColoc over PICCOLO is that it could be run on a larger fraction of simulated datasets (97% versus 39%).

**Figure 3:**
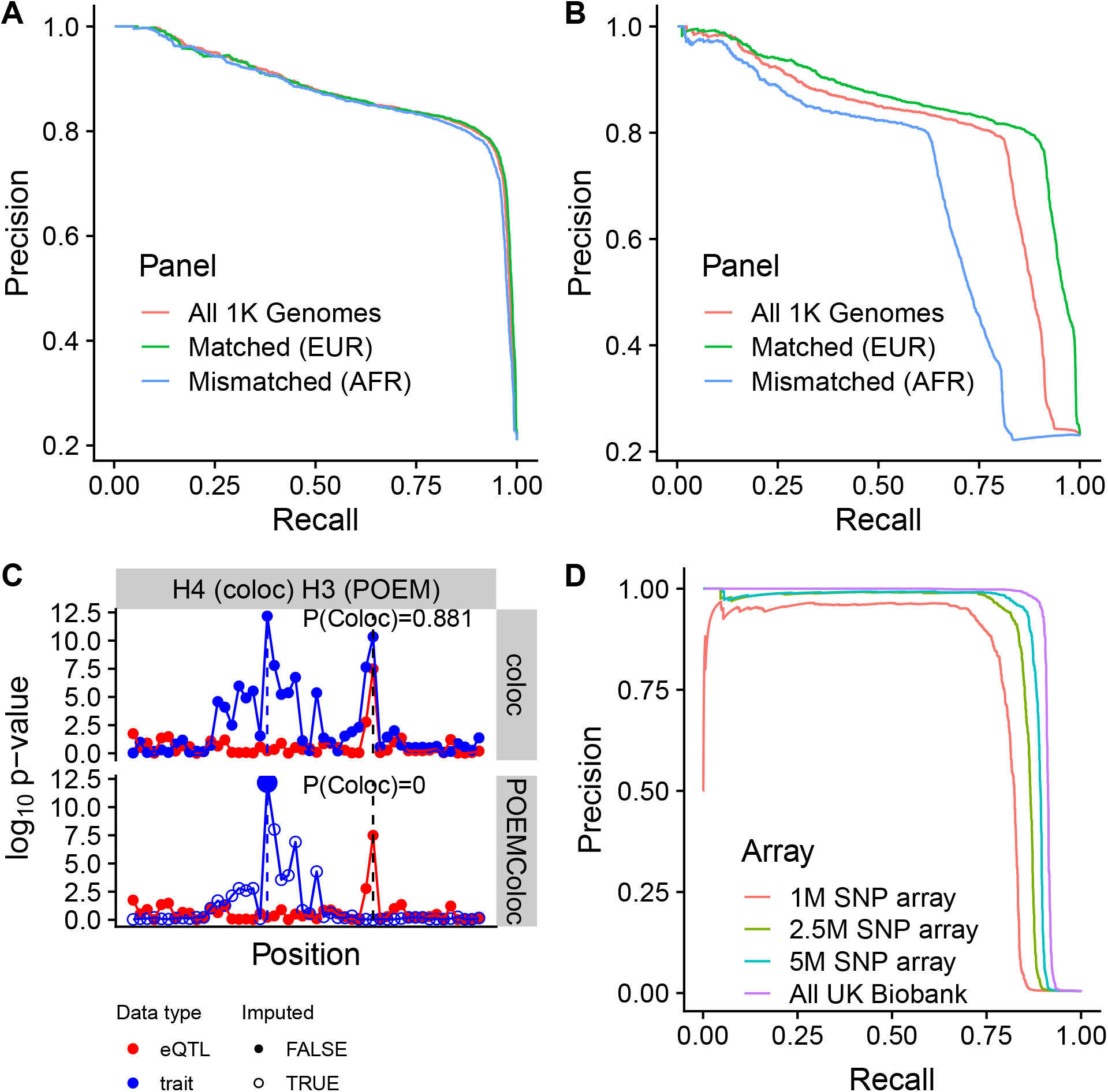
Scenarios that reduce POEMColoc performance A: POEMColoc-1 sensitivity to mismatched ancestry panels. Performance at identifying colocalization (*H*_4_) in datasets simulated under *H*_0_ – *H*_4_. B: POEMColoc-2 sensitivity to mismatched ancestry panels. Performance at identifying colocalization (*H*_4_) in datasets simulated under *H*_0_ – *H*_4_. C: An example of a dataset in which there are multiple independent GWAS signals. Only the second peak is colocalized with the eQTL, so POEMColoc does not detect it. D: Precision-recall curves for predicting colocalization using coloc (*P*(*H*_4_) > 0.9) in the UK Biobank where the POEMColoc top SNP was restricted to SNPs on different sized Illumina array genotyping subsets (1M - OmniExpressExom8; 2.5M - HumanOmni2.5Exome; 5M - HumanOmni5Exome)

### 4.3 POEMColoc caveats

#### 4.3.1 Effect of reference panel ancestry

We next compared performance of POEMColoc when the ancestry of the reference panel was mismatched to the GWAS study ancestry. We can see that when GWAS data are simulated using British ancestry genotypes, performance of POEMColoc is better using individuals from the 1000 genomes European superpopulation as a reference panel than when using all 1000 genomes individuals or a mismatched superpopulation. Differences between ancestry panels are minor given that the GWAS top SNP exists in ancestry panel and when only one of the two datasets requires imputation (Figure 3A). The largest difference comes from the proportion of top SNPs that can be found in the panel (and therefore the proportion of datasets to which POEMColoc can be applied). While only 3% of GWAS top SNPs from the simulation were missing from the 1000 Genomes European samples, 12% were missing from the 1000 Genomes African samples. When both datasets require imputation, effects are more substantial (Figure 3B). In simulation, mismatched ancestry generally reduces recall more than precision using a typical cutoff value of 0.9 (Table S2). Overall, we find that a well-matched ancestry panel improves performance, but that the method will be usable, but conservative in the presence of mismatched ancestry.

#### 4.3.2 Multiple independent associations in region

While we find that across all tested regions, the correlation between colocalization probabilities is very high, there were notable exceptions. In particular, there is a population of tests which has high (> 0.9) probability of colocalization using full summary statistics, but that has a low < 0.1) probability of colocalization using POEMColoc (a false negative population). This accounts for 9% of datasets with colocalization posterior probability greater than 0.9 using coloc with full summary statistics. Within the population of false negatives, we noticed many in which there appear to be two peaks. Figure 3C shows an illustrative example. The lower GWAS peak appears to have a colocalized eQTL peak, but the GWAS peak containing the top SNP does not appear colocalized. Indeed, false negative datasets are enriched for datasets with similar UKBB *p*-value between the GTEx top SNP and UKBB top SNP, but low LD between them, a scenario consistent with multiple independent UKBB causal SNPs (Figure S6A). In this scenario, when we impute new association statistics from only the top UKBB SNP in the region, we only recover association statistics that were in LD with this top peak and which do not have a corresponding eQTL peak. The authors of coloc indicate that colocalization probability is generally based on the strongest association signal, but will be affected by multiple causal variants explaining a similar proportion of variance in the trait, and recommended performing colocalization conditional *p*-values [1]. We find performing a conditional analysis using GCTA-COJO [14] reduces colocalization discrepancies in many datasets (Figure S6B).

#### 4.3.3 Effect when reported SNP comes from limited set of tag variants

We sought to evaluate the performance of the POEMColoc method when only directly genotyped variants were used. To simulate this scenario, we took our original analysis of full UKBB summary statistics and subset them as if one of 3 popular genotyping platforms spanning a range of genomic coverage had been used, now choosing only the top “directly genotyped” variant as the input to POEMColoc. We find that the POEMColoc method tends to miss colocalized signals rather than identify false colocalizations in this analysis and we find that performance suffers most when the set of directly genotyped SNPs is small (Figure 3D, Figure S3).

### 4.4 Application to full GWAS catalog

Having confidence that the POEMColoc method accurately approximated the full coloc method in real UKBB data and that it recovered true colocalized signals generated from simulation, we sought to apply the POEMColoc method to GWAS results for which full summary statistics are not available. Figure 4A summarizes exclusion criteria, cohort ancestry, and study design among analyzed associations. In total, we assessed 47049 reported GWAS associations with *p*-value ≤ 5×10^-8^ for colocalization. Note that associations excluded due to missing information were usually from difficult to parse sample size and design and in most cases should be possible to analyze using POEMColoc after manual review of the entry or linked PubMed article. Additional associations were excluded due not being able to locate the associated SNP in our reference panel. Colocalizations with posterior probability greater than 0.9 are provided in Supplementary Dataset 1.

GWAS Catalog associations meeting inclusion criteria were tested for colocalization using GTEx eQTL in 48 tissues using the POEMColoc method. Our analysis detected 151247 colocalized association-cis gene-tissue triplets, or 0.2% of those tested. Figure 3B shows the number of colocalized genes detected per association. Using cutoff 0.9, 44% of associations have at least one detected colocalization with a GTEx eQTL in some tissue, and 22% have colocalizations for more than one gene. Given there are one or more colocalized genes, the median number of tissues involved is 2, though much larger numbers of tissues are also common.

**Figure 4:**
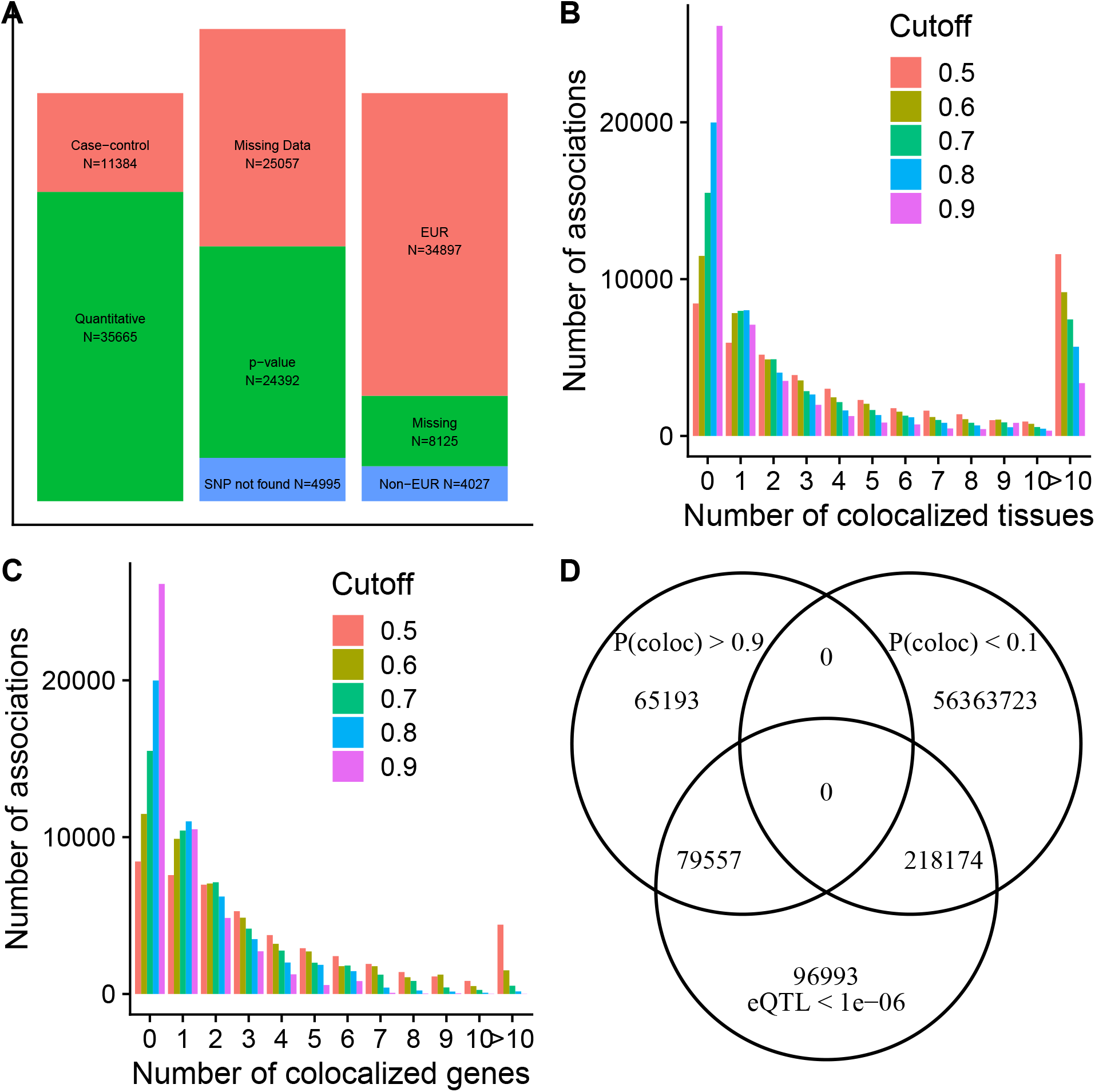
A: Number of GWAS Catalog associations by design, superpopulation, and inclusion criteria. B: Number of colocalized tissues per GWAS Catalog association. C: Number of colocalized genes per GWAS Catalog association. D: Overlap between colocalized association-gene-tissue trios (*P*(*H*_4_) > 0.9) and eQTL significance (*p* < 10^-6^) of the reported GWAS Catalog top SNP.

#### 4.4.1 Tissue Enrichment

To evaluate the biological relevance of POEMColoc output, we assess whether colocalizations are enriched in disease-related tissues. Figure 5A shows tissue enrichments for traits with both large numbers of colocalizations and significant enrichment in some tissue. GTEx tissues have been collapsed into tissue groups as provided by GTEx. These enrichments are largely biologically interpretable, for example enrichment of blood count phenotypes in whole blood, lipid, cholesterol and protein measurements in the liver, and hypothyroidism in the thyroid. Supplementary Table S5 gives a complete list of enrichments detected. We also provide a comparison to enrichments that would be detected using eQTL *p*-values and show that some biologically interpretable enrichments (e.g. QT interval in the heart left ventricle) are only detected using POEMColoc. Because of the large-scale nature of this analysis and because we wish to reduce the impact of human biases in evaluating the method, we quantify disease-related tissues using an automated approach in the MeSH vocabulary. Significantly enriched tissue-trait pairs using POEMColoc are more similar than those that are not enriched (Figure 5B).

**Figure 5:**
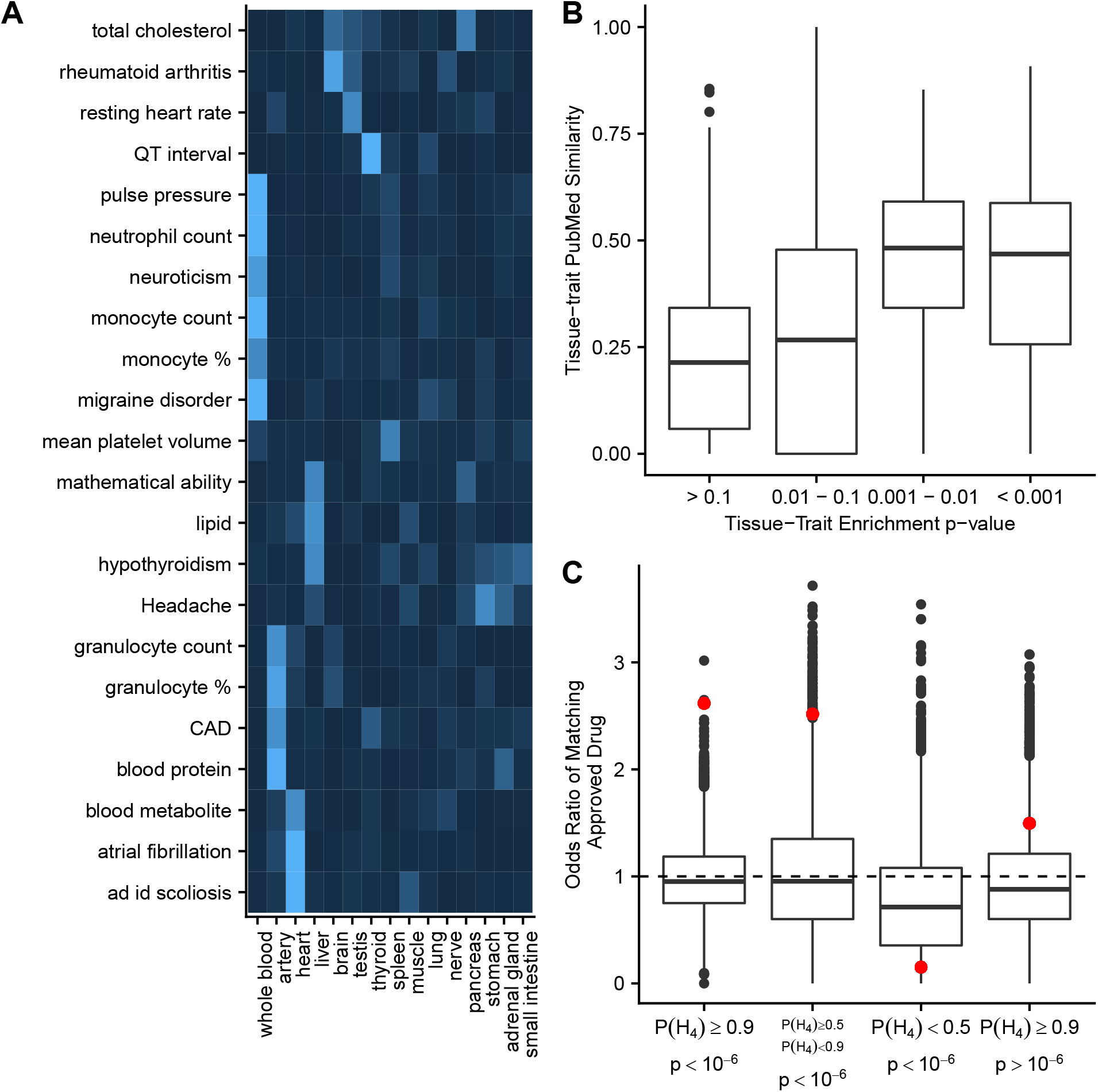
A: Heat map of tissue enrichments for select traits with the largest enrichments. Brighter blue indicates more significant enrichment. CAD = coronary artery disease, ad id scoliosis = adolescent idiopathic scoliosis, granulocyte % = granulocyte percentage of myeloid white cells, monocyte % = monocyte percentage of leukocytes B: Average tissue-trait similarity for enriched pairs by enrichment *p*-value. Similarities are computed from cooccurance in PubMed articles. C: Odds ratio of a gene-trait pair matching an approved drug mechanism by eQTL *p*-value of the GWAS top SNP and colocalization posterior probability P(*H*_4_). Odds ratios are computed relative to cis gene-trait pairs with both eQTL *p*-value > 10^-6^ and colocalization posterior probability less than 0.9.7

#### 4.4.2 Ability to predict approved drug mechanisms

We ask whether colocalized associations are enriched for matches to approved drug mechanisms, an independent source of evidence for target involvement in disease using methodology and supplementary datasets from [15] and [12]. Table S6 shows matches to approved drug mechanisms from colocalized GWAS Catalog eQTL. We find significant enrichment for both significant top SNP eQTL and colocalized eQTL (Figure 5C). The largest enrichment is found when there is both an eQTL significant at 10^-6^ and high to intermediate posterior colocalization probability. There are few matches to approved drug mechanisms for significant eQTL with low *P*(*H*_4_)(< 0.5). In part, this may be because these eQTL are less likely to be coding, and if coding, are less likely to be drug targets than those with high or intermediate colocalization posterior probability (Figure S7). Upon examination of Table S6 we see many closely related drug mechanisms are supported by colocalization evidence (e.g. PPARA for Hypercholesterolemia and Hyperlipidemias). In order to account for this, we counted the number of distinct targets with approved drug mechanisms, and find significantly higher counts than expected by chance (35 targets, *p* < 0.0001).

## 5 Discussion

We report a new method for performing colocalization analysis when only the summary statistics at a lead SNP are reported. We show that the method provides a close approximation to the popular coloc method in application to real association summary statistics from the UK Biobank and GTEx projects.

The most common disagreement between the POEMColoc method and the coloc method were coloc gave a high probability of colocalization and POEMColoc a low probability. We find that this “false negative” population is enriched for regions where there is evidence of multiple causal variants in the GWAS trait. When multiple causal variants are present, imputed summary statistics used by the POEMColoc method will only be accurate for variants linked to the top SNP.The ‘false negative” population is relatively smaller in our simulated datasets, but occurs more often in mismatched ancestry panels (Table S4).

Our method uses a population reference panel to obtain LD estimates and from these, we impute missing summary statistics in the region. As LD can differ between populations of different ancestries, we wondered how the method would perform if there was a mismatch between reference panel ancestry and the original GWAS ancestry. Our simulations show that mismatched ancestry has a minor effect on POEMColoc-1 and while the best performance is achieved when GWAS and reference panel ancestries are matched, overall ability to correctly assign causal configurations in simulated data remains high when using completely mismatched ancestry panels (e.g., all European vs all African). We observed a more substantial decay in performance of POEMColoc-2 and POEMColoc-1 when the top SNP reported in the GWAS catalog comes from a limited subset of directly typed variants.

We show that colocalized gene/trait pairs are enriched in tissues relevant to the underlying GWAS trait. Some of these enrichments seem intuitive and confirm colocalizations inferred by the POEMColoc method can recover known relationships between traits and relevant tissues. Other trait-tissue enrichments are not as intuitive but are supported by known disease biology. Total cholesterol, low density lipoprotein, and lipid levels are enriched for eQTL colocalizations in the liver, a major site of both production and metabolism of cholesterol and LDL. Migraine and headaches are most enriched for colocalizations with artery eQTL and several lines of evidence including the ability of vasoactive substances to induce migraine, the effective treatment of migraines with drugs acting in the vascular system, and increased comorbidity of migraines and cardiovascular diseases support a causal role of the vascular system on migraine [16]. Overall, we find that significant enrichments in colocalizations between traits and tissues is predictive of the co-occurrence rates of the same traits and tissues in the PubMed literature.

Drug targets with genetic support for their therapeutic hypothesis are more likely to result in approvals than drug targets without genetic support [15, 12]. We recently demonstrated that this increase in approval probability seems to depend on the type and quality of genetic support with genetic support derived from Mendelian associations and GWAS associations with a clear causal gene having a stronger effect than GWAS associations where the causal gene was more uncertain [12]. We tested whether colocalized gene-trait pairs were enriched among approved target-indication pairs. Indeed, we find that gene-trait pairs that have evidence of colocalization and evidence of an eQTL association at the trait associated locus are enriched by approximately 3-fold in approved target-indication pairs. This enrichment is not found in gene-trait pairs that have evidence of an eQTL at the trait associated locus but do not show evidence of colocalization with the trait. Thus, evidence of colocalization from the POEMColoc method may be useful for identifying drug targets that are more likely to succeed in clinical development.

## 6 Acknowledgements

We thank Samantha Lent for help using GCTA-cojo and Xiuwen Zheng for help using the SeqArray R package.

