## Supplementary Figures and Tables for "Estimating colocalization probability from limited summary statistics"

### S1 POEMColoc maximizes likelihood under approximate normal distribution

Assume a region with $p$ SNPs. Assume a single causal SNP and let $c$ be its index. Let $\lambda$ be the expected value of the $Z$ statistic at the causal SNP. The log likelihood under the approximate normal distribution is

$$l(\lambda, c|Z) = \frac{1}{2}\log|2\pi\Sigma| - \frac{1}{2}\left(Z - \lambda r_c\right)' \Sigma^{-1}\left(Z - \lambda r_c\right) \tag{1}$$

Absorbing constant terms with respect to the parameters

$$l(\lambda, c|Z) = \lambda r_c' \Sigma^{-1} Z - \frac{1}{2}\lambda^2 r_c' \Sigma^{-1} r_c = \lambda Z_c - \frac{1}{2}\lambda^2 \tag{2}$$

The simplification is due to $r_c$ being a row of $\Sigma$. This is maximized at $\lambda = Z_c$ and $c = \text{argmax} Z_i$. The maximizing choice of $c$ may not be unique, and the model may not be identifiable if two markers have $r = 1$.

Using these plugin estimates for $c$ and $\lambda$ we can impute $Z_{2,mis}$ with its conditional expectation given $Z_{obs}$.

$$\mu_{mis|obs} = Z_{obs} r_{mis,obs} \tag{3}$$

### S2 Tissue enrichment computations

Let $C_{ijk}$ be 1 if trait $i$ is found to be colocalized with an eQTL of gene $j$ in tissue $k$ and 0 otherwise. We compute an enrichment odds ratio for tissue $I$ and trait $K$ and associated $p$-value from Fisher's exact test using the following $2 \times 2$ table.

$$\begin{matrix} \sum_j C_{IjK} & \sum_j \sum_{k \neq K} C_{Ijk} \\ \sum_j \sum_{i \neq I} C_{ijK} & \sum_j \sum_{k \neq K} \sum_{i \neq I} C_{ijk} \end{matrix}$$

Conditioning on colocalization when performing the enrichment analysis serves to mitigate the effect of confounding variables such as different colocalization probabilities across genes, traits and tissues. To motivate this computation consider the following model

$$p(C_{i,j,k} = 1) = \mu \alpha_i \beta_j \gamma_k \delta_{ik} \eta_{ij} \tau_{jk}$$

If $\delta_{ik} = 1$ for all $i, k$ and $\tau_{j,k} = 1$ (no interactions between colocalization in tissue and colocalization in trait, and no interactions between colocalization in gene and colocalization in trait) for all $j, k$, expected cell counts are

$$\begin{matrix} \mu \alpha_I \gamma_K \sum_j \beta_j \eta_{Ij} & \mu \alpha_I \sum_j \beta_j \eta_{Ij} \sum_{k \neq K} \gamma_k \\ \mu \gamma_K \sum_{i \neq I} \sum_j \beta_j \alpha_i \eta_{ij} & \mu \sum_{i \neq I} \sum_j \beta_j \alpha_i \eta_{ij} \sum_{k \neq K} \gamma_k \end{matrix}$$

leading asymptotically to an odds ratio of

$$\frac{\mu \alpha_I \gamma_K \sum_j \beta_j \eta_{Ij} \mu \sum_{i \neq I} \sum_j \beta_j \alpha_i \eta_{ij} \sum_{k \neq K} \gamma_k}{\mu \alpha_I \sum_j \beta_j \eta_{Ij} \sum_{k \neq K} \gamma_k \mu \gamma_K \sum_{i \neq I} \sum_j \beta_j \alpha_i \eta_{ij}} = 1$$

### S3 Follow up on UK Biobank analysis using GTCA-COJO

The most common difference between POEMColoc and coloc applied to the UK Biobank was that a minority of datasets with high estimated colocalization probability using coloc applied to full summary statistics had low estimated colocalization probability using POEMColoc. We defined a "false negative" colocalization to be one in which coloc estimated the posterior probability of colocalization greater than 0.9, but POEMColoc estimated the posterior probability of colocalization be less than 0.1 and subjected them to further investigation. When looking at datasets "false negative" colocalization using POEMColoc, many appeared to have two separate peaks in the full dataset, but only one peak in the imputed dataset (Figure 3C). One of the peaks was colocalized with the eQTL, but it was not the highest. To address this, we ran GCTA-COJO [1] on the full summary statistics from UK Biobank to detect multiple causal SNPs within a 2 Mb window surrounding each top SNP from the UK Biobank using a $p$-value cutoff of $5 \times 10^{-8}$ and colinearity parameter 0.8. A subsample of 5,000 individuals from the UK Biobank was used as a reference panel. When multiple SNPs were detected, we ran a conditional analysis for the top SNP conditional on all other SNPs. In some cases, the top SNP was not selected in the conditional analysis. If a SNP with LD at least 0.9 to it was, we used this in its place, otherwise the dataset was excluded from the conditional colocalization analysis. We recomputed colocalization probabilities using the conditional summary statistics as input to POEMColoc in place of the raw summary statistics. As a point of comparison, we also looked at how the same analysis changes the results for true positive or true negative datasets (defined by both POEMColoc and coloc having colocalization probabilities greater than 0.9 and less than 0.1 respectively).

### S4 Precision-recall curve confidence intervals

Area under the precision-recall curve (AUCPR) was computed using the `precrec` `R` package [2]. Bootstrap confidence intervals were generated using `boot R` [3] for individual methods and pairwise differences in AUCPR. Following recommendations [4], we used a stratified bootstrap.

Table S1: Performance of methods in predicting coloc at cutoff 0.9 in the UK Biobank and predicting causal causal variants in simulation. Area under precision recall curve (AUCPR) and 95% bootstrap confidence interval. All confidence intervals for pairwise differences in AUCPR exclude zero (from bootstrap resampling of pairwise differences), except for the difference between PICCOLO and POEMColoc-2 on simulated datasets. % missing gives the percent of datasets that were not analyzed due to missing SNP information.

| method | AUCPR UKBB | AUCPR Simulation | Missing % UKBB | Missing % Simulation |
|---|---|---|---:|---:|
| POEMColoc-1 | 0.9 (0.84,0.95) | 0.9 (0.89,0.91) | 5.3 | 3.3 |
| PICCOLO | 0.74 (0.66,0.82) | 0.87 (0.86,0.89) | 26.2 | 60.9 |
| eQTL p-value | 0.32 (0.25,0.38) | 0.81 (0.79,0.82) | NA | NA |
| POEMColoc-2 | 0.8 (0.73,0.87) | 0.88 (0.87,0.89) | 5.9 | 6.6 |
| coloc | NA | 0.9 (0.89,0.91) | NA | NA |

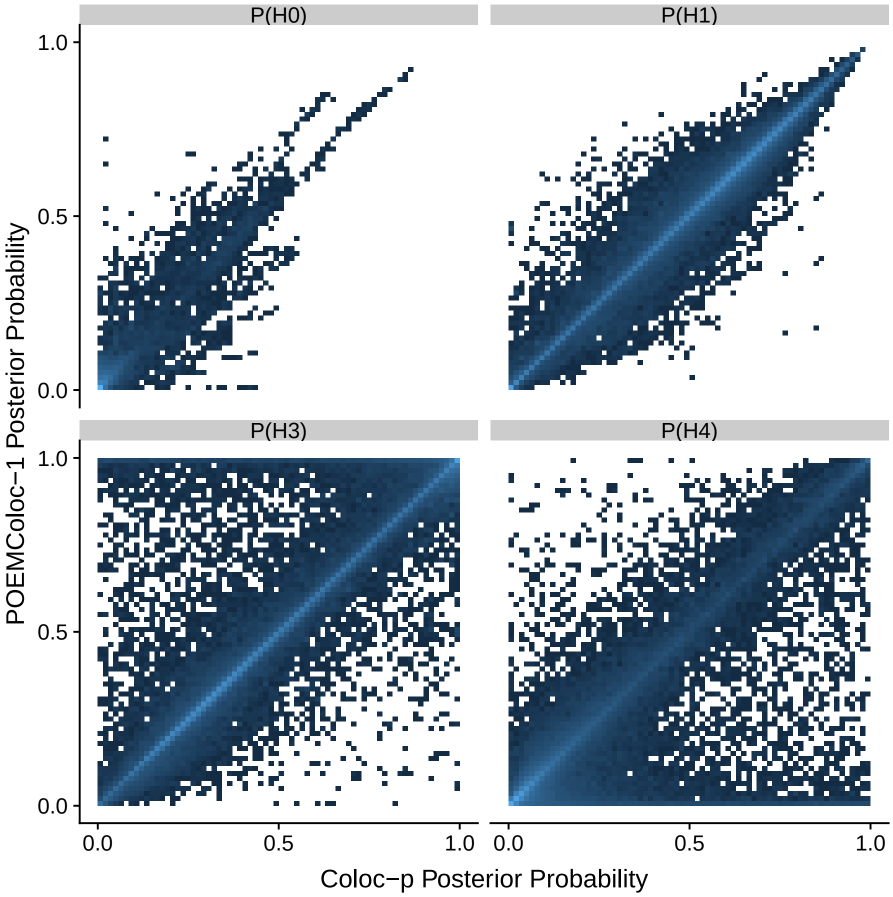


Figure S1: Comparison of hypothesis posterior probability estimates using POEMColoc-1 method to coloc using GTEx eQTL regression coefficients and UK Biobank *p*-value summary statistics as input (coloc-p)

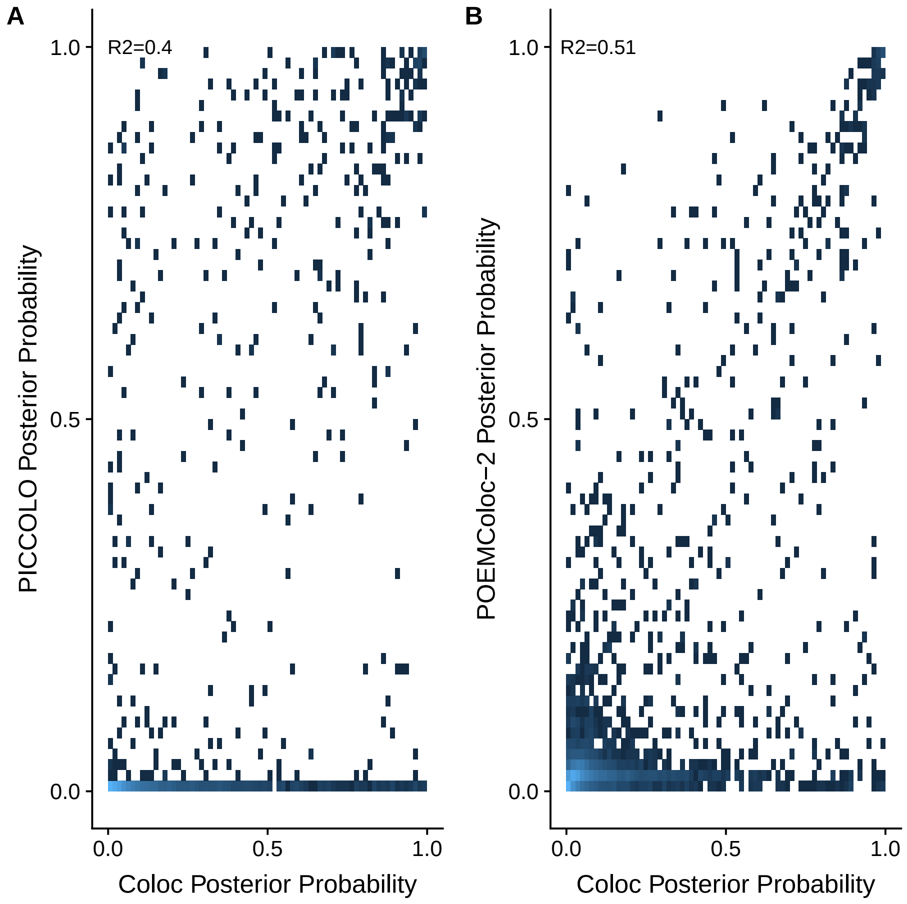


Figure S2: A: Comparison of colocalization posterior probability estimates between PICCOLO and coloc for GTEx eQTL and 100 UK Biobank phenotypes. B: Comparison of colocalization posterior probability estimates between POEMColoc-2 and coloc for GTEx eQTL and the same 100 UK Biobank phenotypes.

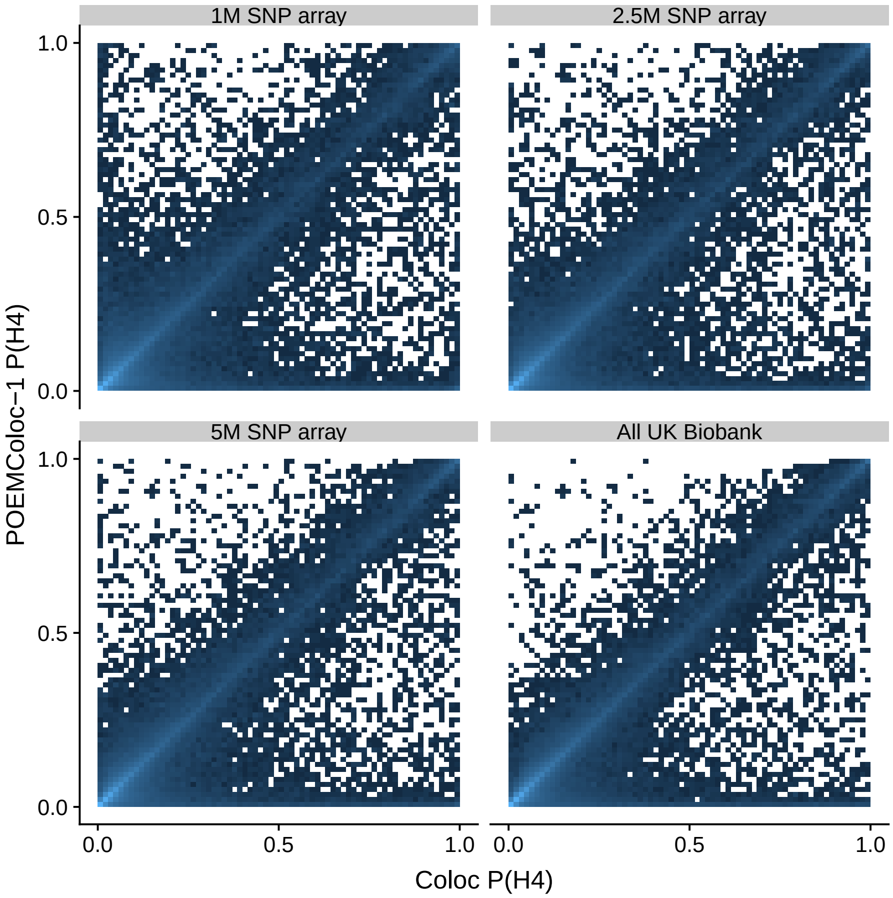


Figure S3: POEMColoc-1 colocalization posterior probability versus colocalization probability estimated using coloc on the UK Biobank and GTEx whole blood eQTL. Comparison of hypothesis posterior probability estimates for coloc and POEMColoc-1 where the POEMColoc top SNP was restricted to SNPs on different sized Illumina array genotyping subsets (1M - OmniExpressExom8; 2.5M - HumanOmni2.5Exome; 5M - HumanOmni5Exome)

Table S2: Sensitivity of POEMColoc performance to ancestry panel misspecification in simulation study. Genotypes from UK Biobank European ancestry were used in the simulation, so 1000 genomes EUR is expected to be the best-matching panel. Missing % is the percent of simulated datasets that could not be analyzed due to the top SNP not being in the panel. Recall and precision are computed using a cutoff value of 0.9.

| Method | Panel | AUCPR | Missing % | Recall | Precision |
|---|---|---|---:|---:|---:|
| POEMColoc-1 | EUR | 0.88 (0.87,0.89) | 3.3 | 0.68 | 0.83 |
| POEMColoc-1 | AFR | 0.88 (0.87,0.89) | 12.4 | 0.66 | 0.84 |
| POEMColoc-1 | All 1K Genomes | 0.88 (0.87,0.89) | 3.2 | 0.67 | 0.84 |
| POEMColoc-1 | AMR | 0.88 (0.87,0.89) | 4.1 | 0.67 | 0.83 |
| POEMColoc-1 | SAS | 0.88 (0.87,0.89) | 7.3 | 0.68 | 0.83 |
| POEMColoc-1 | EAS | 0.86 (0.85,0.87) | 28.9 | 0.65 | 0.82 |
| POEMColoc-2 | EUR | 0.86 (0.85,0.87) | 6.6 | 0.67 | 0.83 |
| POEMColoc-2 | AFR | 0.72 (0.7,0.73) | 22.5 | 0.36 | 0.85 |
| POEMColoc-2 | All 1K Genomes | 0.81 (0.8,0.83) | 6.3 | 0.50 | 0.85 |
| POEMColoc-2 | AMR | 0.84 (0.83,0.85) | 8.1 | 0.59 | 0.84 |
| POEMColoc-2 | SAS | 0.84 (0.82,0.85) | 13.7 | 0.61 | 0.83 |
| POEMColoc-2 | EAS | 0.8 (0.78,0.81) | 47.0 | 0.51 | 0.82 |

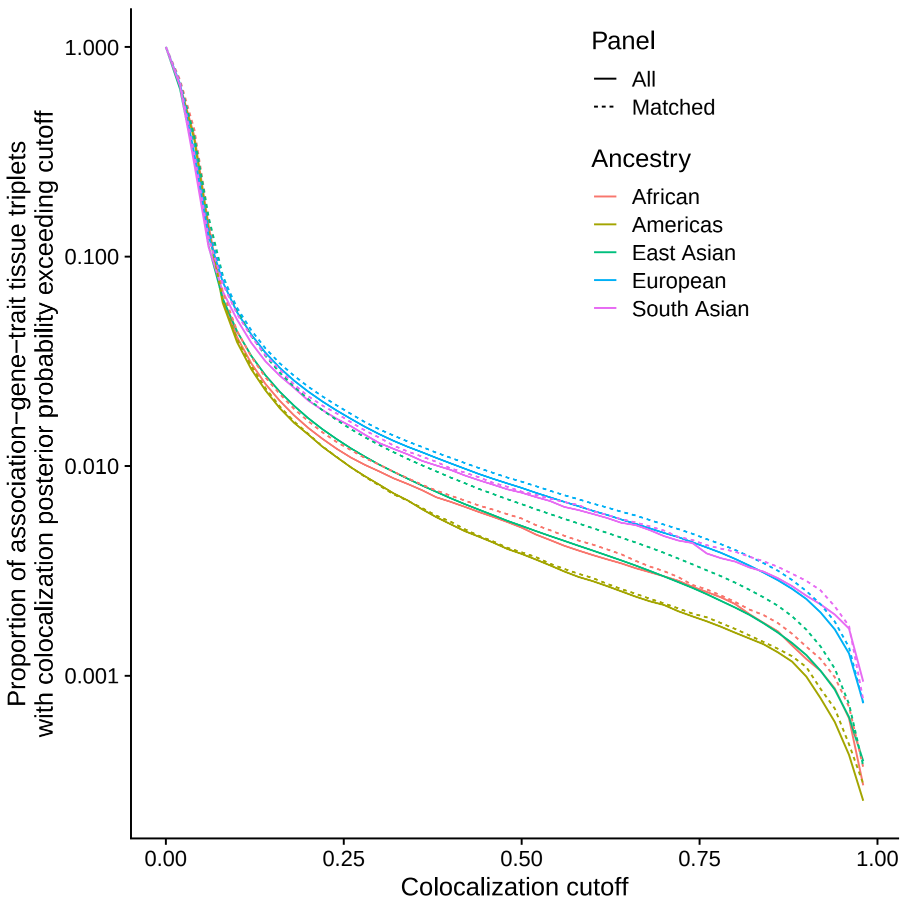


Figure S4: Proportion of association-gene-trait tissue triplet showing evidence for colocalization by colocalization cutoff. Colors correspond to study ancestry and solid and dashed lines to whether the imputation panel is matched ancestry or all 1000 Genomes related individuals.

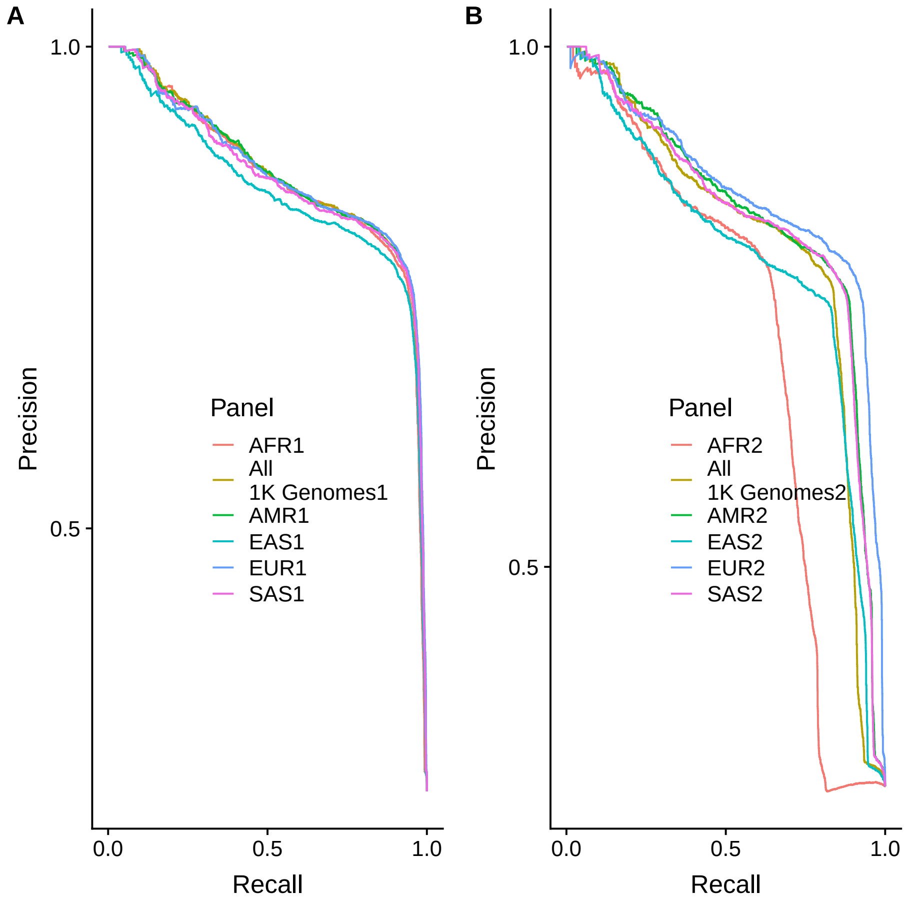


Figure S5: Simulation performance of POEMColoc using different ancestry panels. Left: POEMColoc with single top SNP dataset. Right: POEMColoc with two top SNP datasets. Ancestry panels are taken from unrelated Phase 3 1000 Genomes superpopulation individuals (EUR, EAS, SAS, AFR, AMR) or all 1000 Genomes unrelated individuals (All). EUR is expected to have the best performance because the genotypes used in simulation were from British ancestry individuals from UK Biobank.

Table S3: Proportion p of SNP-trait pairs with true positive and false negative colocalization that have multiple associations detected by COJO. n is the number analyzed, excluding those where the top SNP was not recognized in the reference panel.

| class | n | p |
|---|---|---|
| FN | 212 | 0.52 |
| TN | 349 | 0.42 |
| TP | 378 | 0.39 |

Table S4: Proportion of coloc true positives that are false negatives using POEMColoc-1 in simulation. A coloc true positive is defined as a dataset simulated under colocalization with coloc posterior probability $> 0.9$. A POEMColoc-1 false negative is a dataset simulated under colocalization with POEMColoc-1 posterior probability less than 0.1. These are defined in order to be comparable to the UK Biobank analysis.

| Panel | POEM_FN |
|---|---|
| African | 0.011 |
| All | 0.009 |
| Americas | 0.009 |
| East Asian | 0.007 |
| European | 0.004 |
| South Asian | 0.006 |

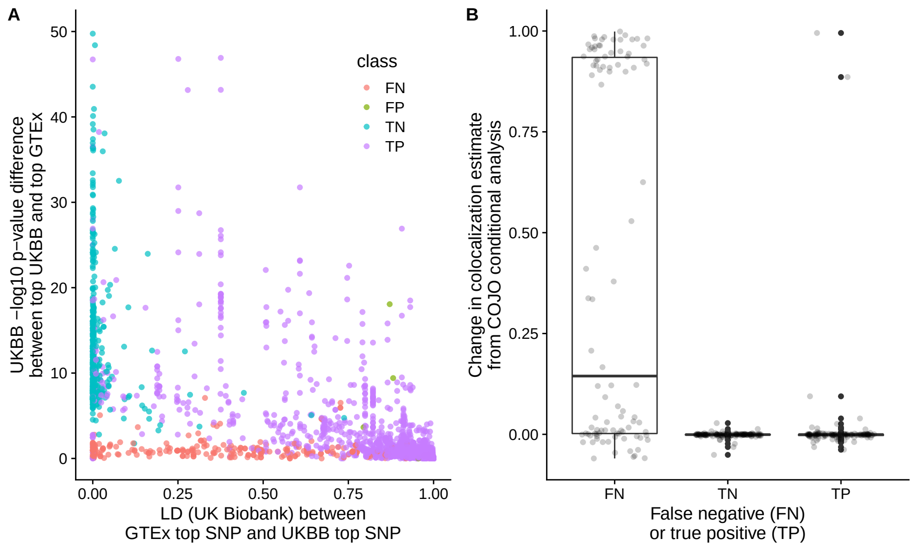


Figure S6: A: LD between GWAS top SNP and GTEx top SNP (x) and difference in GTEx p-value between GTEx top SNP and GWAS top SNP (y) discriminate true positive, false negative, and false positive colocalizations estimated using POEMColoc colocalization of UK Biobank associations and GTEx whole blood expression. Truth is defined to be positive if colocalization probability estimated from the full data is $> 0.9$ and negative if $< 0.1$, and positive and negatives from POEMColoc are defined with the same cutoffs. Points in which either method gave a colocalization estimate between 0.1 and 0.9 are excluded. y-axis truncated at 50. B: Reanalyzing datasets using COJO conditional summary statistics for top SNP conditional on all other associations disproportionately effects performance in false negative datasets compared to true positives.

Table S5: Top tissue trait enrichment -log10 p-values detected by POEMColoc-1 and eQTL p-value approach. POEMColoc-1 uses a colocalization cutoff of 0.9. eQTL p-value cutoff of $\sim 10^{-12}$ was chosen such that the total number of gene-trait links between the two methods was the same. Because of the arbitrary choice of thresholds, using the same number of top-ranked links ensures we are using thresholds of similar stringency. All enrichments having p-value 0.05 or lower using either method, and at least 3 genes with significant eQTL or colocalizations for the trait in the tissue are shown.

| trait | tissue | POEM | pvalue | Similarity |
|---|---|---|---|---|
| atrial fibrillation | heart_atrial_appenda | 12.00 | 4.02 | 0.91 |
| granulocyte percentage of myel | whole_blood | 9.85 | 10.13 | 0.52 |
| atrial fibrillation | heart_left_ventricle | 9.69 | 3.85 | 0.62 |
| granulocyte count | whole_blood | 8.10 | 5.50 | 0.52 |
| neutrophil count;eosinophil co | whole_blood | 7.71 | 5.99 | 0.48 |
| monocyte percentage of leukocy | whole_blood | 7.19 | 8.25 | 0.45 |
| neutrophil count;basophil coun | whole_blood | 6.24 | 5.55 | 0.48 |
| pulse pressure measurement | artery_aorta | 5.67 | 3.31 | 0.65 |
| neutrophil count | whole_blood | 5.54 | 5.90 | 0.48 |
| QT interval | heart_left_ventricle | 5.50 | 1.01 | 0.73 |
| C-C motif chemokine 5 measurem | cells_ebv-transforme | 5.38 | 0.00 | NA |
| migraine without aura | artery_tibial | 5.36 | 4.00 | 0.00 |
| hypothyroidism | thyroid | 5.33 | 1.60 | 0.81 |
| ejection fraction measurement | heart_left_ventricle | 5.11 | 3.14 | 0.74 |
| left ventricular systolic func | heart_left_ventricle | 5.11 | 3.14 | 0.78 |
| fractional shortening | heart_left_ventricle | 5.11 | 3.14 | 0.79 |
| migraine disorder | artery_aorta | 4.59 | 1.72 | 0.23 |
| glucose measurement | thyroid | 4.55 | 2.41 | 0.40 |
| coronary artery disease | artery_aorta | 4.45 | 3.61 | 0.54 |
| monocyte count | whole_blood | 4.38 | 3.74 | 0.45 |
| lipid measurement | liver | 4.10 | 1.09 | 0.58 |
| Behcet's syndrome | whole_blood | 3.98 | 1.89 | 0.27 |
| age at menarche;hip bone miner | heart_atrial_appenda | 3.87 | 1.35 | 0.00 |
| Agouti-related protein measure | thyroid | 3.86 | 3.11 | NA |
| mean platelet volume | whole_blood | 3.67 | 0.89 | 0.00 |
| systemic scleroderma;systemic | cells_transformed_fi | 3.62 | 3.88 | 0.65 |
| apolipoprotein A 1 measurement | liver | 3.60 | 4.88 | 0.62 |
| granulins measurement | liver | 3.60 | 4.88 | NA |
| adolescent idiopathic scoliosi | testis | 3.60 | 0.15 | 0.20 |
| total cholesterol measurement | liver | 3.52 | 1.10 | 0.61 |
| repulsive guidance molecule A | liver | 3.51 | 4.88 | 0.35 |
| mammographic density measureme | adipose_subcutaneous | 3.38 | 1.35 | 0.00 |

Table S5: Top tissue trait enrichment -log10 p-values detected by POEMColoc-1 and eQTL p-value approach. POEMColoc-1 uses a colocalization cutoff of 0.9. eQTL p-value cutoff of $\sim 10^{-12}$ was chosen such that the total number of gene-trait links between the two methods was the same. Because of the arbitrary choice of thresholds, using the same number of top-ranked links ensures we are using thresholds of similar stringency. All enrichments having p-value 0.05 or lower using either method, and at least 3 genes with significant eQTL or colocalizations for the trait in the tissue are shown. *(continued)*

| trait | tissue | POEM | pvalue | Similarity |
|---|---|---:|---:|---:|
| femoral neck bone mineral dens | artery_tibial | 3.37 | 1.26 | 0.27 |
| blood metabolite measurement | stomach | 3.31 | 0.33 | NA |
| myeloid white cell count | whole_blood | 3.28 | 5.55 | 0.20 |
| migraine without aura | artery_aorta | 3.28 | 0.00 | 0.32 |
| granulins measurement | minor_salivary_gland | 3.23 | 0.00 | NA |
| rheumatoid arthritis | spleen | 3.22 | 1.90 | 0.33 |
| systemic scleroderma;systemic | spleen | 3.20 | 0.95 | 0.51 |
| alcohol drinking | testis | 3.20 | 0.00 | 0.22 |
| Takayasu arteritis | whole_blood | 3.19 | 0.42 | 0.18 |
| migraine disorder | artery_tibial | 3.18 | 3.04 | 0.35 |
| body mass index;hematocrit;glu | liver | 3.05 | 2.13 | NA |
| serum alanine aminotransferase | liver | 2.94 | 0.94 | NA |
| Abdominal Aortic Aneurysm | liver | 2.78 | 4.88 | 0.15 |
| alcoholic pancreatitis | pancreas | 2.58 | 2.93 | 0.82 |
| breast size | adipose_subcutaneous | 2.50 | 3.46 | 0.54 |
| lipoprotein-associated phospho | liver | 2.38 | 1.37 | NA |
| complement factor H measuremen | liver | 2.29 | 3.17 | NA |
| catalase measurement | liver | 2.15 | 3.17 | NA |
| interleukin 18 receptor 1 meas | whole_blood | 1.99 | 3.63 | NA |
| plasminogen activator inhibito | liver | 1.97 | 4.34 | NA |
| C-reactive protein measurement | liver | 1.75 | 3.05 | 0.50 |
| coronary artery disease | artery_tibial | 1.62 | 3.04 | 0.49 |
| diastolic blood pressure;smoki | cells_transformed_fi | 1.50 | 5.24 | NA |
| serum gamma-glutamyl transfera | liver | 1.43 | 4.50 | NA |
| prostate specific antigen meas | adrenal_gland | 0.85 | 3.48 | NA |
| systolic blood pressure;smokin | cells_transformed_fi | 0.70 | 4.34 | NA |
| coffee consumption;cups of cof | thyroid | 0.48 | 5.07 | 0.19 |
| progranulin measurement | liver | 0.34 | 3.82 | NA |
| vital capacity | brain_putamen_basal_ | 0.28 | 3.02 | 0.00 |
| systolic blood pressure;alcoho | cells_transformed_fi | 0.26 | 3.35 | NA |
| QRS complex;QRS amplitude | thyroid | 0.16 | 8.28 | 0.22 |
| bipolar disorder;schizophrenia | esophagus_mucosa | 0.00 | 4.16 | 0.00 |

Table S5: Top tissue trait enrichment -log10 p-values detected by POEMColoc-1 and eQTL p-value approach. POEMColoc-1 uses a colocalization cutoff of 0.9. eQTL p-value cutoff of $\sim 10^{-12}$ was chosen such that the total number of gene-trait links between the two methods was the same. Because of the arbitrary choice of thresholds, using the same number of top-ranked links ensures we are using thresholds of similar stringency. All enrichments having p-value 0.05 or lower using either method, and at least 3 genes with significant eQTL or colocalizations for the trait in the tissue are shown. *(continued)*

| trait | tissue | POEM | pvalue | Similarity |
|---|---|---|---|---|
| response to bronchodilator;for | muscle_skeletal | 0.00 | 3.55 | NA |
| response to bronchodilator;FEV | muscle_skeletal | 0.00 | 3.55 | NA |
| smoking status measurement;uri | colon_transverse | 0.00 | 3.45 | NA |
| intracranial volume measuremen | brain_amygdala | 0.00 | 3.12 | 0.25 |
| coronary heart disease;lipopro | liver | NA | 4.88 | NA |
| coronary heart disease;acute c | liver | NA | 4.88 | NA |
| t-tau measurement | thyroid | NA | 3.10 | NA |

Table S6: Approved drug mechanisms with colocalization.

| indication | symbol | Trait | sim |
|---|---|---|---|
| Psoriasis | ITGAL | Crohn's disease;psoriasis;ulce | 1.00 |
| Psoriasis | PDE4A | Crohn's disease;psoriasis;ulce | 1.00 |
| Crohn Disease | PTGS2 | Crohn's disease | 1.00 |
| Psoriasis | IL23A | psoriasis | 1.00 |
| Schizophrenia | DRD2 | schizophrenia | 1.00 |
| Depression | DRD2 | depressive symptom measurement | 1.00 |
| Arthritis, Rheumatoi | IL6R | rheumatoid arthritis | 1.00 |
| Psoriasis | IL6R | Crohn's disease;psoriasis;ulce | 1.00 |
| Arthritis, Juvenile | IL6R | systemic juvenile idiopathic a | 1.00 |
| Asthma | ADORA1 | asthma | 1.00 |
| Atrial Fibrillation | SCN5A | atrial fibrillation | 1.00 |
| Colitis, Ulcerative | PTGS2 | Crohn's disease | 0.90 |
| Factor X Deficiency | F10 | prothrombin time measurement | 0.88 |
| Arthritis, Psoriatic | PDE4A | Crohn's disease;psoriasis;ulce | 0.87 |
| Infertility, Female | LHCGR | polycystic ovary syndrome | 0.84 |
| Hyperlipidemias | LPL | triglyceride measurement | 0.83 |
| Inflammatory Bowel D | PTGS2 | Crohn's disease | 0.83 |
| Hypercholesterolemia | NPC1L1 | total cholesterol measurement | 0.82 |
| Hypercholesterolemia | HMGCR | total cholesterol measurement | 0.82 |
| Hypercholesterolemia | PCSK9 | total cholesterol measurement | 0.82 |
| Hypercholesterolemia | PPARA | total cholesterol measurement | 0.82 |

Table S6: Approved drug mechanisms with colocalization. *(continued)*

| indication | symbol | Trait | sim |
|---|---|---|---|
| Hypercholesterolemia | LPL | triglyceride measurement;high | 0.82 |
| Psychotic Disorders | DRD2 | schizophrenia | 0.82 |
| Hyperlipoproteinemia | LPL | triglyceride measurement | 0.81 |
| Tachycardia, Ventric | SCN5A | PR interval | 0.81 |
| Disseminated Intrava | F10 | prothrombin time measurement | 0.81 |
| Disseminated Intrava | F2 | prothrombin time measurement | 0.81 |
| Arthritis, Juvenile | IL1R1 | Crohn's disease;psoriasis;ulce | 0.80 |
| Thyroid Neoplasms | FGFR3 | bladder carcinoma | 0.80 |
| Arrhythmias, Cardiac | KCNH2 | QT interval | 0.80 |
| Arrhythmias, Cardiac | SCN5A | PR interval | 0.80 |
| Hypophosphatasia | ALPL | vitamin B6 measurement | 0.80 |
| Arthritis | PDE4A | Crohn's disease;psoriasis;ulce | 0.80 |
| Bipolar Disorder | DRD2 | schizophrenia | 0.79 |
| Arthritis, Rheumatoi | IL1R1 | Crohn's disease;psoriasis;ulce | 0.79 |
| Hemophilia B | F7 | prothrombin time measurement | 0.79 |
| Hemophilia B | F10 | prothrombin time measurement | 0.79 |
| Hemophilia B | F2 | prothrombin time measurement | 0.79 |
| Hyperlipidemias | NPC1L1 | total cholesterol measurement | 0.79 |
| Hyperlipidemias | HMGCR | total cholesterol measurement | 0.79 |
| Hyperlipidemias | PCSK9 | total cholesterol measurement | 0.79 |
| Hyperlipidemias | PPARA | total cholesterol measurement | 0.79 |
| Hyperlipoproteinemia | NPC1L1 | total cholesterol measurement | 0.79 |
| Hyperlipoproteinemia | APOB | total cholesterol measurement | 0.79 |
| Hyperlipoproteinemia | HMGCR | total cholesterol measurement | 0.79 |
| Hyperlipoproteinemia | PCSK9 | total cholesterol measurement | 0.79 |
| Ventricular Fibrilla | KCNH2 | QT interval | 0.79 |
| Ventricular Fibrilla | SCN5A | PR interval | 0.79 |
| Multiple Sclerosis, | IL2RA | multiple sclerosis | 0.78 |
| Depressive Disorder, | DRD2 | neuroticism measurement | 0.78 |
| Dermatitis, Atopic | PDE4A | Crohn's disease;psoriasis;ulce | 0.78 |
| Hypertension | ADRB1 | systolic blood pressure | 0.77 |
| Hypertension | AGT | systolic blood pressure | 0.77 |
| Hypertension | ACE | diastolic blood pressure | 0.77 |
| Hypertension | NOS3 | diastolic blood pressure | 0.77 |
| Thyroid Neoplasms | TSHR | Graves disease | 0.76 |
| Hypotension | AGT | mean arterial pressure | 0.76 |
| Bronchitis, Chronic | ADORA1 | asthma | 0.75 |
| Atherosclerosis | LPL | triglyceride measurement;high | 0.75 |
| Neutropenia | CSF3 | neutrophil count | 0.75 |
| Depressive Disorder, | MTNR1B | conduct disorder | 0.75 |

Table S6: Approved drug mechanisms with colocalization. *(continued)*

| indication | symbol | Trait | sim |
|---|---|---|---|
| Chest Pain | SCN5A | PR interval | 0.74 |
| Hypersensitivity | ADORA1 | asthma | 0.74 |
| Constriction, Pathol | PDE4A | Crohn's disease;psoriasis;ulce | 0.74 |
| Endocrine System Dis | TPO | hypothyroidism | 0.74 |
| Atrial Fibrillation | KCNH2 | QT interval | 0.74 |
| Hypertriglyceridemia | HMGCR | total cholesterol measurement | 0.73 |
| Hypertriglyceridemia | PPARA | total cholesterol measurement | 0.73 |
| Myelodysplastic Synd | DNMT1 | myeloid white cell count | 0.73 |
| Dermatitis, Atopic | IL4R | eosinophil count | 0.73 |
| Hemophilia A | F7 | prothrombin time measurement | 0.73 |
| Hemophilia A | F10 | prothrombin time measurement | 0.73 |
| Hemophilia A | F2 | prothrombin time measurement | 0.73 |
| Hypogonadism | LHCGR | polycystic ovary syndrome | 0.73 |
| Leukemia, Myeloid, A | FLT3 | granulocyte percentage of myel | 0.73 |
| Leukemia, Myeloid, A | DNMT1 | granulocyte count | 0.73 |
| Hemorrhage | F7 | prothrombin time measurement | 0.73 |
| Hemorrhage | F10 | prothrombin time measurement | 0.73 |
| Hemorrhage | F2 | prothrombin time measurement | 0.73 |
| Pulmonary Disease, C | ADORA1 | asthma | 0.73 |
| Atherosclerosis | HMGCR | total cholesterol measurement | 0.71 |
| Atherosclerosis | PPARA | total cholesterol measurement | 0.71 |
| Breast Neoplasms | ERBB2 | age at menopause | 0.71 |
| Diabetes Mellitus, T | LPL | triglyceride measurement;high | 0.71 |
| Kidney Neoplasms | FGFR3 | bladder carcinoma | 0.70 |
| Leukemia, Myelogenou | DNMT1 | myeloid white cell count | 0.70 |

[4] K. Boyd, K. H. Eng, and C. D. Page, "Area under the precision-recall curve: point estimates and confidence intervals," in *Joint European confer-*

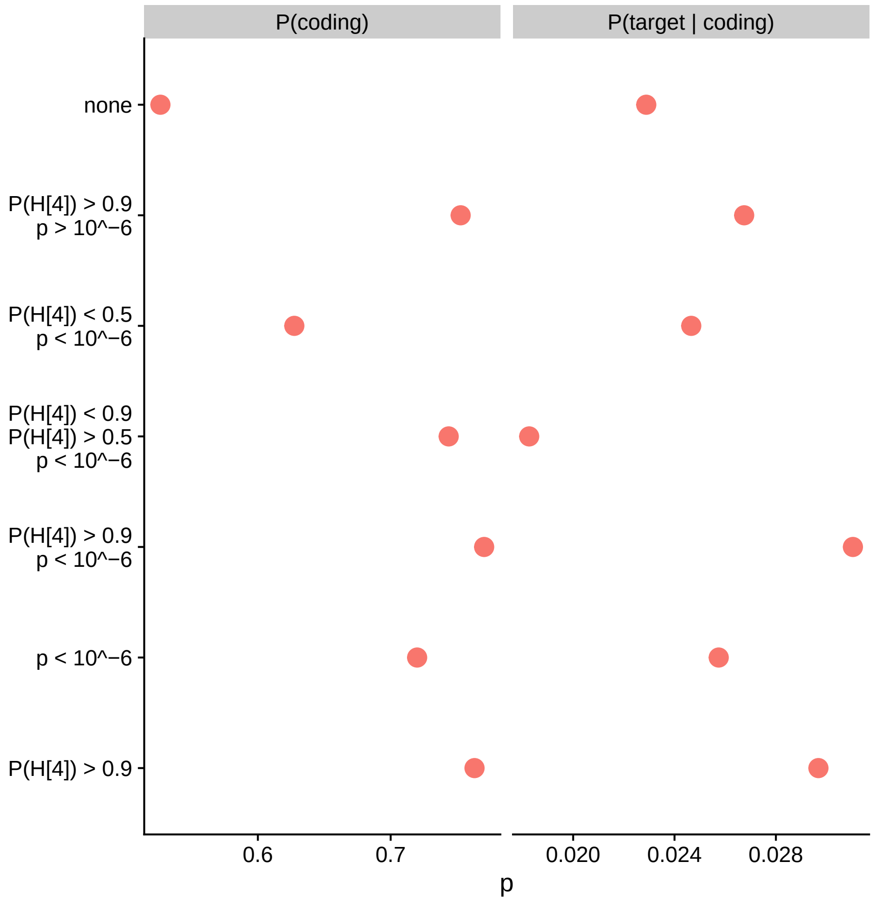


Figure S7: Proportion of gene-trait pairs eQTL $p$-value of the GWAS top SNP and colocalization posterior probability $\mathrm{P}(H_4)$ meeting cutoffs that involve coding genes (A) and approved drug targets conditional on being coding (B). xMHC genes were excluded from both the numerator and the denominator of B because they are unavailable in approved drug target dataset.

*ence on machine learning and knowledge discovery in databases*, pp. 451–466, Springer, 2013.
